# Hydrogen Bond Networks and Hydrophobic Effects in the Amyloid *β*_30–35_ Chain in Water: A Molecular Dynamics Study

**DOI:** 10.1101/090092

**Authors:** KwangHyok Jong, Luca Grisanti, Ali Hassanali

## Abstract

We study the conformational landscape of the C-terminal fragment of the Amyloid protein A*β*_30–35_ in water using well-tempered metadynamics simulations and find that it resembles an intrinsically disordered protein. The conformational fluctuations of the protein are facilitated by a collective reorganization of both protein and water hydrogen bond networks, combined with electrostatic interactions between termini as well as hydrophobic interactions of the side chains. The stabilization of hydrophobic interactions in one of the conformers involves a collective collapse of the sidechains along with a squeeze out of water sandwiched in between. The charged N and C termini play a critical role in stabilizing different types of protein conformations including those involving contact ion salt-bridges as well as solvent mediated interactions of the termini and amide backbone. We examine this by probing the distribution of directed water wires forming the hydrogen bond network enveloping the polypeptide. Water wires and their fluctuations form an integral part of structural signature of the protein conformation.

## 1 Introduction

There is currently an active effort from both experimental and theoretical fronts to understand the physical and chemical process underlying of protein fibril formation as well as of the early stage of aggregation^1–24^. These fibrils are three-dimensional architectures resulting from the aggregation of misfolded proteins. One of the most studied fibrils in this regard, are those that develop from the Amyloid *β*(A*β*) protein, a sequence made up of 39-43 amino acids. Besides serving as excellent model systems to understand physical and chemical processes in a biological context, the Amyloid fibrils have been implicated in neurodegenerative diseases such as Alzheimers and Parkinsons.^25^ Apart from the eventual plaques that form, there is a growing appreciation that oligomers such as dimers and trimers can also play a critical role in the pathology of Alzheimers disease.^26^ There is thus a lot of interest to understand the conformational heterogeneity of A*β* since the structural disorder that features it during the early stages of aggregation may have important implications on its subsequent dynamical evolution.^27^ Besides the A*β* protein fibrils, there have been numerous experimental studies showing that fibril-like structures can result from the aggregation of different types of poly-peptide chains typically made up of hydrophobic amino-acids.^28–32^ Similar to A*β*, these fibrils are characterized by a dense network of hydrogen bonds between the polar backbone making beta-sheet secondary structures. The exact details of how hydrogen bonding and hydrophobic interactions couple with each other remains an open question. A very recent experimental study for example, used solid-state NMR to show that the abundance of hydrophobic amino acids in A*β*1-42 results in a dense packing of alkylic side chains in the plain perpendicular to the fibril axis.^33^ Much less is known however, about how polar and non-polar interactions within the protein and the surrounding solvent couple with each other before aggregation has even started.

Recently, some of us have been involved in trying to understand a rather peculiar and anomalous experimental observation, namely that amyloid fibrils are capable of fluorescing in the absence of aromatic residues.^34^ Using state-of-the-art first principles simulations on small model amyloid crystals, we find that salt-bridges between the N and C termini are characterized by strong hydrogen bonds where proton transfer leads to the formation of both zwitterionic and non-zwitterionic states in the fibril. This feature is tuned by the surrounding hydrogen bond network involving the proximity of water and hydrophobic amino acids.^35^ An obvious limitation of the tools deployed in this earlier study is that both the model systems and time-scales did not allow us to explore the larger scale conformational fluctuations associated with the hydrogen bond network.

Providing a detailed microscopic description of both the structural landscape and molecular interactions for the monomer with the surrounding solvent bath is key to understanding how subsequent aggregation proceeds. There have thus been numerous theoretical studies examining the conformational fluctuations of both the A*β* monomer as well as smaller segments of it.^36–43^ In many of these studies, the model monomer peptides are terminated with two methyl groups and this by construction, hinders the possible formation of strong hydrogen bonds between the N-C termini and possibly between the termini and the backbone.

In this work, we use classical molecular dynamics (MD) simulations to explore the protein and water networks of the hydrated C-terminal hydrophobic peptide of A*β*(A*β*_30–35_). This chain formed a segment of one of the model crystals we studied in our earlier work with first principle methods.^34^ Furthermore, Liu et al.^40^ have shown that A*β*_30–35_ plays a critical role in the aggregation process by including short anti-parallel strands in the surrounding residues, which in turn could promote the fibril formation of full-length A*β*. Using well-tempered metadynamics, we explore the free energy landscape of this chain in its zwitterionic, NH_3_^+^.**AIIGLM**·COO^−^(NC) form. We also perform simulations of a variant of NC where the termini ends are instead capped with methyl groups CH_3_CONH· **AIIGLM**·CONHCH_3_(MET) to examine the importance of the termini. The change in the termini interactions leads to significant changes in the conformational landscape of NC and MET. We find that the disorder in the conformational landscape of NC is driven by a diversity of different interactions such as pure electrostatic interaction between termini, polar interactions of the backbone, van-der-Waals interactions of the hydrophobic side chains and also hydrogen bonds between the termini and the backbone.

We also investigate the topological properties of the hydrogen bond network surrounding the peptide as probed through the reorganization of water wires connecting N-donor (N−H and NH_3_^+^) and carbonyl (C=O) groups. Similar types of analysis have been conducted to understand the network structures of bulk water, aqueous solutions and also in understanding the mechanisms of proton transfer in water.^44–46^ In addition, Thirumalai and co-workers have also shown the formation of single-file water wires between two sheets of the yeast prion protein although these have been investigated mostly in a qualitative manner.^47,48^ Our analysis reveal subtle differences such as the shortening or lengthening of the wires connecting different parts of the peptide during the conformational fluctuations, providing new insights into the coupling of protein and water motions for this system. The changes in these wires often include those involving the N and C termini in NC and provide a rationale for understanding the changes in the structural disorder when one moves to the MET system.

The paper is organized as follows. We begin in Section 2 with details of the computational protocols employed in this work. We then move on in Section 3.1 to discuss our results on the free energy landscapes of the NC and MET systems as revealed by our metadynamics simulations. Here, we also discuss the coupling of polar and non-polar interactions in stabilizing the disordered structures observed in NC. In Section 3.2, we discuss our analysis of the reorganization of hydrogen bond networks around the peptide. Finally, we end in Section 4 with some conclusions and possible future directions of our work.

## 2 Methods

Below we highlight the molecular dynamics simulation details including a brief overview of the theory behind metadynamics and also how our collective variables were chosen for the metadynamics simulations.

### 2.1 Simulation Details

The system NC consists of the 6 amino acid sequence AIIGLM. The starting structure for this hexapeptide was extracted from one of the 8 chains forming the model amyloid crystal structure with PDB code 2Y3J.^49^ For the NC system, the termini were terminated with NH_3_^+^ and COO^−^ groups corresponding to pH=7 solvent conditions. The MET structure is obtained by modifying the termini of NC by capping the two ends with methyl-amide and acetyl groups. NC and MET are solvated with 7052 and 7098 water molecules respectively. The two systems were subsequently run for 20 ns in the NPT ensemble using the Berendsen barostat^50^ at 1 atm and using a time constant of 0.5 ps. The final cubic box lengths used for NC and MET were then 6.0 nm. All simulations were thermostated at 300K with a Nosé-Hoover thermostat^51,52^ using a time constant of 0.1 ps. A non-bonded pair list was produced using a cut-off radius of 1.4 nm. The short-range non-bonded pairwise interactions were evaluated by using a shifted Lennard-Jones potential with a cut-off length at 0.9 nm while the long-range electrostatic interactions were handled using the Particle Mesh Ewald-Switch (PME-switch) method^53,54^ with a Coulomb switching cut-off at 1.2 nm. A long range dispersion correction was applied to both energy and pressure for the van-der-Waals cutoff. No bond-constraints were imposed in the simulation and hence a smaller time-step of 0.5 fs is employed for the Verlet integration.

For all our simulations, the OPLS-AA force field and TIP4P water model were employed. In a recent study, Smith and co-workers did a systematic comparison of a wide variety of forcefields including OPLS-AA, AMBER, CHARMM and GROMOS on the conformational landscape of A*β*21-30 in water compared to available experimental data.^55^ Their conclusion was that the OPLS-AA forcefield suppressed the formation of helical structures consistent with the experiments and hence recommended the use of either OPLS or GROMOS for their system. Since we are also doing simulations on a segment of the amyloid protein, we decided to conduct our simulations with OPLS/AA and TIP4P.

### 2.2 Free Energy Calculations

Free energy surfaces were explored using well-tempered metadynamics (WT-MetaD)^56^ simulations. Here, we briefly summarize the key theoretical concepts behind the methodology. A history-dependent bias potential, 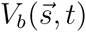 (where the quantity 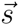 is the vector of collective variables), is introduced to enhance the sampling of the free energy surface in the basis of a predefined set of collective variables. 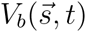 is a sum of Gaussian hills with the width 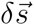 and the height w, centered at the values of the collective variables that have already been visited and deposited at the time interval *τ*_*G*_,

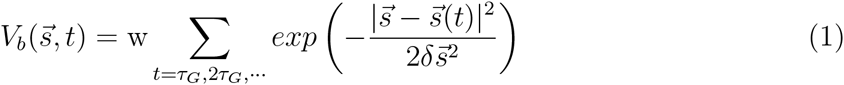

Unlike standard metadynamics, the height of Gaussian hills added is modified according to the relationship

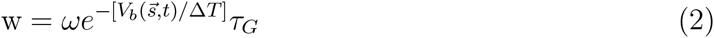

where *ω* is the initial bias deposition rate with dimension of energy rate and Δ*T* is a tunable temperature-like parameter that controls how quickly w reduces as the wells are filled. The parameters, *ω* and Δ*T*, are chosen to achieve the best efficiency. The free energy can then be computed with the following expression

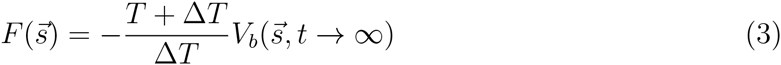

In this work, we have biased two collective variables based on our analysis of some relatively short unbiased simulations of the NC system. Over the course of a 80ns MD simulation, we observed a single event where the N and C termini came into close proximity forming a strong salt-bridge for a couple of nanoseconds before dissociating away from each other. Therefore we have choosen the end-to-end distance between and the N and C termini, *d*_*ee*_, as one of the CVs for the WT-MetaD. As we will see later in the manuscript, backbone hydrogen bonding and sidechain hydrophobic packing also plays an important role in the structural disorder we observe in the peptide conformations. In order to enhance the fluctuations in these coordinates, we also biased the radius of gyration, *R*_*g*_, as another CV as has been done in previous studies.^21,22^

As we will see later, while we have only biased two CVs namely, *d*_*ee*_ and *R*_*g*_, we also construct free energy surfaces in other variables that are not biased. In order to do this, re-weighted free energy profiles along unbiased coordinates were constructed with a recent reweighting algorithm.^57^ Some examples of collective variables that we found to be important to understand the structural disorder in NC and MET include, the extent of hydrogen bonding between the backbone amide groups, van-der-Waals packing of the hydrophobic side chains and the exposure of both the backbone and the side-chains to the surrounding water. In all these cases, contact maps defined by a switching function were used. See SI (Sect. SI-1) for more details on the details of the functional form of these variables as well as the parameters used to define them.

Besides the quantities involving the reorganization of the protein backbone and side chain, We also examined various topological properties like the water networks connecting different parts of the protein. In particular, we examined directed water wires connecting donor and acceptor groups of the peptide. Donor groups include the N-terminus and amide N−H bonds of the backbone, while the C-terminus and the carbonyl C=O groups are acceptor groups. Oxygens of water molecules (O_w_) act as both donors and acceptors. All the donors and acceptors are treated as vertices on a graph with edges between vertices. The vertices include the nitrogen and oxygen atoms of the protein and the oxygen atoms of water molecules. If two vertices *v*_*i*_ and *v*_*j*_ are connected by an edge *e*_*ij*_, they are said to be adjacent. The edges *e*_*ij*_ encode information about the hydrogen bonds. In this work, we assign a weight to edges leading to a weighted directed adjacency matrix A=A(G), which is an *M* × *M* matrix with M being the number of vertices. There are broadly two criterion to define a hydrogen bond, one based on geometry and another based on energetics. In this work, we use the former and define the weights of the edges based on a combination of the distance and angle of the hydrogen bond.^58^ More specifically, the elements of adjacency matrix, *A*_*ij*_, for the weighted graph are defined as

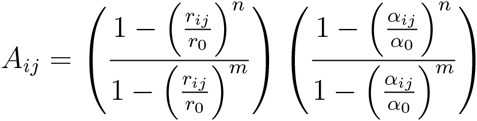

where *r*_*ij*_ is the distance between donor and acceptor of H-bond and *α*_*ij*_ is the angle H–D–A, *r*_0_, *α*_0_, m, and n are parameters of the edge weight. Note that in the expression above for *A*_*ij*_, *n* > *m*. The directed water wires are determined by Djisktra’s algorithm^59^ which explores the shortest path of the network. In order to take the advantage of this algorithm, we give small weights into the strong hbond wires and therefore set exponent n with lager value than m. The parameters *r*_0_ and *α*_0_, as thresholds of switching function, for O_w_ −O_w_, N−O_w_ and O–O_w_ hydrogen bonds, were set by the values corresponding to the first minima position of distribution function of both the distance between donor and acceptor and the angle H−D−A. Specific value for the above parameters are listed in the SI, at Sect. SI-2.

All the molecular dynamics simulations were performed using the GROMACS 4.6.7^60^ package and the metadynamics calculations were conducted using the PLUMED2.1^61^ plugin. For both WT-MetaD simulations of NC and MET, the bias factors, *γ* = (*T* +Δ*T*)/Δ*T*, were set to 10 and Gaussian functions were deposited every 1ps with an initial height of 0.5kJ/mol, whereas the widths in *R*_*g*_ and *d*_*ee*_ of Gaussian functions for NC were 0.01nm and 0.01nm and widths in *R*_*g*_ and *d*_*ee*_ for MET were 0.02nm and 0.02nm respectively. These widths were determined from the fluctuations observed during the unbiased simulations of both systems. The simulations of NC and MET chain were run for simulation time of 1.5*μ*s an 1.1*μ*s. The visualization of the structures in this work was done using VMD.^62^

## 3 Results

In this section we move on to characterizing the free energy surfaces (FES) obtained for the NC and MET systems. As we will shortly see, changing the termini has quite a drastic effect on the underlying FES and more particularly, the structural disorder that is observed in the polypeptide chain. We rationalize the origin of this by elucidating the re-organization of both the protein and water hydrogen bond networks.

### 3.1 Free Energy Surfaces

#### 3.1.1 1D-FES

As described earlier, we biased two CVs *d*_*ee*_ and *R*_*g*_ in our WT-MetaD simulations. We begin by illustrating in Figure 1 the 1d-FES along *d*_*ee*_ obtained for the NC and MET systems. This comparison also helps us build our intuition on the underlying interactions that are important for stabilizing different structures we observe. It is clear that just by a cursory visual inspection of the 1d-FES, the two systems NC and MET are very different. In the case of NC, there is a pronounced and narrow minimum at 0.4nm corresponding to the formation of a salt bridge between the N and C terminus, and a much broader basin between 0.8-1.8nm which involve situations where the N and C terminus are far away from each other and hence more solvent exposed. Based on the 1D-FES, there rather large barriers between 7-10kJ/mol to move between these two basins. On the other hand, for MET, the narrow minimum occurs at much larger *d*_*ee*_ distances of about 1.2nm. Approximately 5 kJ/mol above this minimum are two broad and flat basins analagous to what one would see in a typical funnel-like picture of protein folding.^63^

**Figure 1:**
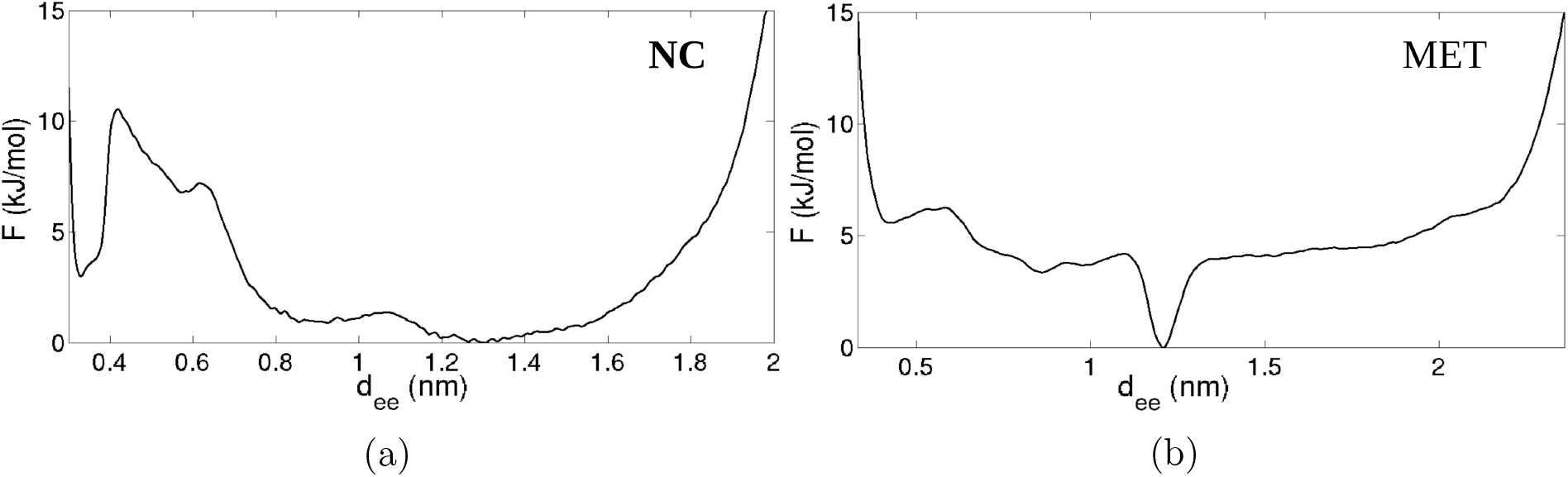
1D FES along *d*_*ee*_ for NC(a) and MET(b)

The differences observed between NC and MET in Figure 1 are not too surprising given that the fluctuations along this coordinate very different driving forces. In the case of NC, there is a strong electrostatic interaction that essentially gets completely unscreened by the solvent in the narrow basin below 0.5nm. For MET, the termini are made up of methyl groups and hence at short *d*_*ee*_ involve much weaker van-der-Waals interactions which simply creates a shallow minimum at around 0.4nm as seen in Figure 1b. This minimum is reminiscent of the contact minimum that occurs in the potential of mean force between two methane molecules in water.^64^ This analysis for both NC and MET integrates out the *R*_*g*_ and hence coordinates that would be involved in characterizing the degree of change in the compactness of the polypeptide chains. We now move on to discuss the higher dimensional FES obtained along both biased and unbiased collective variables in our metadynamics runs.

#### 3.1.2 2D-FES

##### Protein Conformation

We begin by illustrating the 2D FES that is obtained by biasing both the *d*_*ee*_ and *R*_*g*_ CVs for NC in Figure 2. This FES reveals a much richer underlying landscape characterized by structural heterogeneity. The FES features four distinct minima along *d*_*ee*_ and *R*_*g*_ at roughly the following locations: (0.35,0.45), (0.35,0.53), (0.75,0.45) and finally (1.3,0.53). As one might expect based on our earlier observations, the former two at shorter *d*_*ee*_ involve narrower basins due to strong electrostatic forces, while the latter two at larger *d*_*ee*_ are broader owing to the enhanced conformational flexibility. Interestingly, for both situations where there are strong and weak termini interactions in the NC system, there are two basins with smaller and larger radii of gyration which essentially quantifies the extent of compactness or folded character of the system.

**Figure 2:**
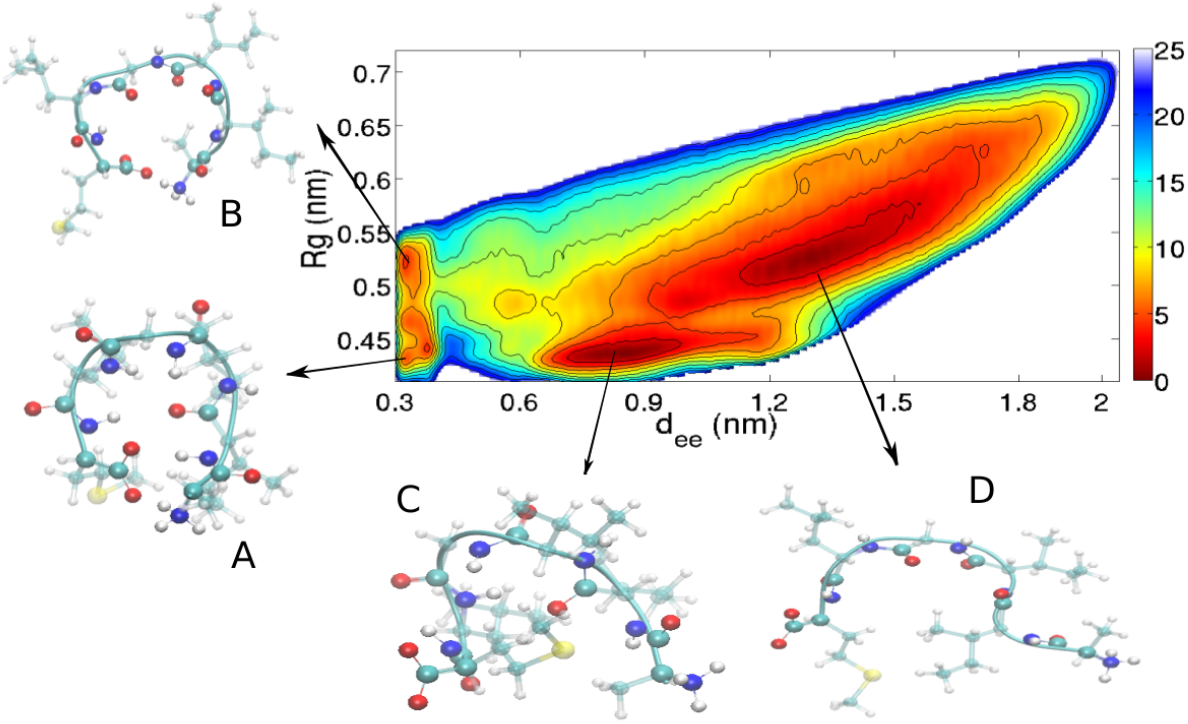
2D FES obtained from 1.5-*μ*s-long WT-MetaD for NC chain. The FES is contoured by 2.5kJ/mol up to 25kJ/mol. The basins and structures depicted on FES correspondent conformational states populated during the simulation time.

In order to aid future discussions in the paper, we label the four basins in Figure 2 A, B, C and D. Also shown in the figure are representative snapshots as a visual guide to the conformational changes that are involved. Within the statistical errors associated with the convergence of our free energies (see SI for details on analysis of the convergence our calculations, Sect. SI-3) all the four states are essentially equally populated. While the NH3+ and COO-groups in both A and B reside very close to each other, the snapshots suggest that both the packing of the backbone and sidechains is quite different. In order to understand these features associated with the states A, B, C and D in more detail, we turn to examining the how changes in the radius of gyration involve the reorganization of both backbone and side-chain packing.

##### Polar and Non-Polar Protein Interactions

To probe deeper into the underlying origins associated with fluctuations along the Rg coordinate, we examined the reweighted FES obtained using two additional unbiased CVs, namely the degree of backbone (BC) and side-chain (SC) contacts as measured by contact maps described earlier in the Methods section. The former quantifies the extent of the hydrogen bonding interactions between the amide bonds as illustrated in the changes observed going from snapshots B to A of Figure 2. The latter reveals changes in the extent of hydrophobic interactions between the side chain groups in the peptide.

The left and right panels of Figure 3 show the 2D-FES obtained from the simulations along *d*_*ee*_ and BC and SC respectively. Overall, we observe a complex coupling of polar and non-polar forces acting simultaneously to stabilize the different conformations. For states A and B which occur for short distances when *d*_*ee*_ is less than 0.5nm, there are three regions along the BC coordinate. We also notice, that the states where the N and C termini are further away from each other (states C and D) are characterized by slightly smaller BC values although the differences are not so drastic indicating that even in the so-called less compact (or extended) states, there are significant hydrogen bonding interactions between the N-H and C=O groups of the backbone. The right panel of Figure 3 illustrates how the hydrophobic side chains contribute to the conformational fluctuations. For short N-C termini distances, there are conformations consisting of high and low SC along the SC coordinate. The changes in the SC parameter in this regime, are quite significant - the value of SC undergoes broad fluctuations and can increase by a factor of 4 between states A and B.

**Figure 3:**
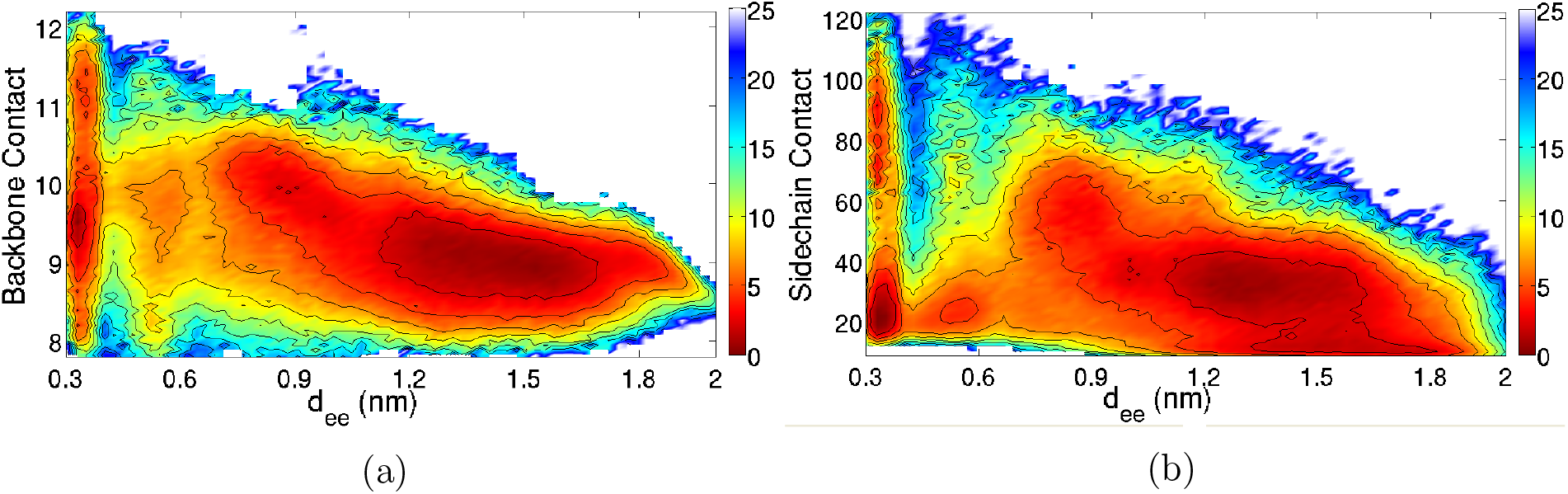
Reweighted 2D FES along *d*_*ee*_ and backbone contact(a) and along *d*_*ee*_ and sidechain contact(b)

The preceding analysis shows the importance of both hydrogen bonding backbone interactions as well as the packing of hydrophobic side chains once the N and C termini are in close proximity. To elucidate how changes in the BC and SC result in changes in the radius of gyration and hence also to resolve better the differences in the origin of states A, B, C, and D in Figure 2, we examined 3D FES’s along *d*_*ee*_, *R*_*g*_, BC and SC as shown in Figure 4. In order to ease the visualization of the 3D FES, we identify three regions of the FES, with different colors: less than 3, 7 and 11 kJ/mol shown in red, yellow and blue respectively. A is characterized by a lower *R*_*g*_ compared to B and based on the 3D FES, the former features slightly larger backbone contacts originating from interactions of the amide dipoles (see Figure 5). In addition, the backbone contacts for state A are quite similar to the extended state C. Interestingly, the 3D-FES shows that the largest BC corresponds to a possible intermediate state that is sampled during the transition between between basins A and B

**Figure 4:**
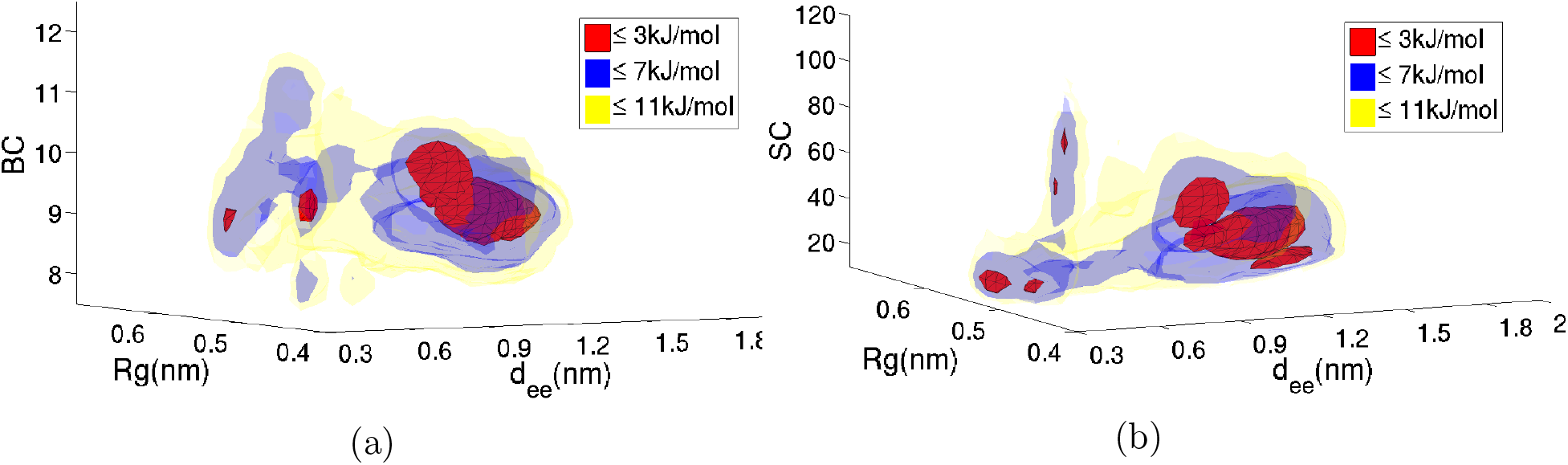
Reweighted 3D FES along *d*_*ee*_, *R*_*g*_ and backbone contact(a) and along *d*_*ee*_, *R*_*g*_ and sidechain contact(b)

**Figure 5:**
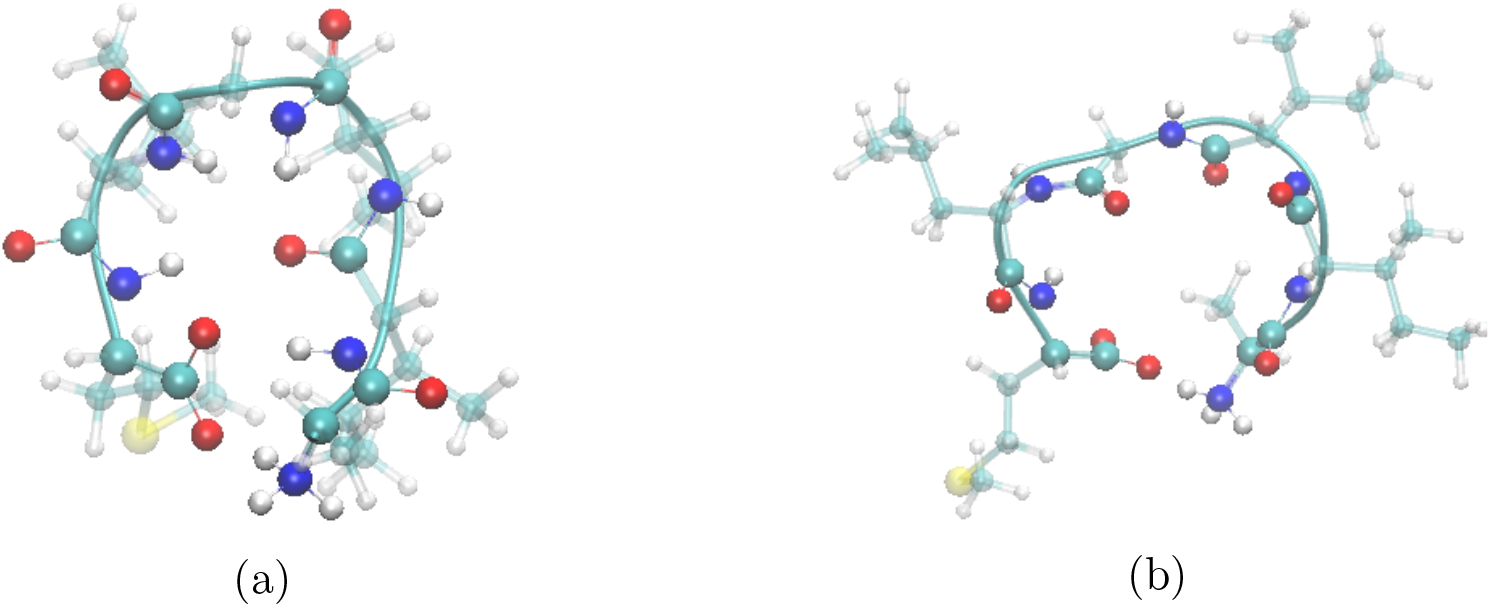
Representative snapshot of basin A(a) and basin B(b)

The right panel of Figure 4 shows the coupling of *d*_*ee*_, *R*_*g*_ and SC and provides better separation of all the various states and also into the origins of the changes of *R*_*g*_. The side chains are all hydrophobic amino acids and hence an increase in their contacts corresponds to the formation of favorable van-der-Waal interactions between alkylic groups. We clearly see here that moving from lower to higher radius of gyration when the N and C termini are in close contact, involves a rather drastic decrease in the extent of side-chain interactions. States C and D on the other hand, are characterized by side-chain contacts sandwiched between A and B. In C, the side chains of amino acids isoleucine and leucine pack closely with each other while in D which has the lowest side-chain packing, involves isoleucine and methionine. Interestingly, moving to state B results in a conformation where all the side chains are solvent exposed. Finally, the transition from B to A involves a collective collapse of all but one of the hydrophobic side chains. This creates a conformation resembling a Janus-like particle with the polar hydrogen bonds of the backbone on one side and the non-polar side chains on the other (see left and right panels of Figure 5). From an energetic standpoint, the creation of this compact Janus-like structure involves a marked decrease of side-chain interaction energies (see SI for details, Figure SI-2 in Sect. SI-4).

The picture developed up to this point, reveals a rich variety of molecular interactions involving both polar hydrogen bonding and hydrophobic interactions for the NC chain. Fluctuations in the backbone and side-chain contacts lead to changes in how compact the peptide chain is and hence on the *R*_*g*_. Although we have focused on the hydrogen bonding interactions of just the amide bonds of the backbone, due to the highly flexible character of the termini, polar interactions *between* the termini and the backbone also play an important role in stabilizing all the various conformations (see SI for further information, Figure SI-3 in Sect. SI-4). These results are also interesting within the context of understanding the aggregation of amyloid fibrils. As indicated in the introduction, the underlying driving forces associated with the formation of the fibrils include both hydrogen bonding of the backbone and burying of hydrophobic side chains.^65^ Our simulations show that these driving forces form an integral part of the conformational fluctuations of our small model peptide.

##### Protein-Water Interactions

It goes without saying that these fluctuations involving the close packing of the hydrogen bond network to stabilize the backbone interactions, as well as the packing of side chains must be intimately coupled to the reorganization of the surrounding solvent. There have been numerous studies in the literature discussing the importance of this coupling.^66–72^ One obvious question that emerges from the preceding analysis regards the changes in the exposure of both the backbone and sidechains to the solvent during the structural transformations along the FES. The left and right panels of Figure 6 show the FES along *d*_*ee*_ and the contact between the backbone/side-chain and water. In constructing this contact map, we focused on water molecules within 3.5 Angstrom of the backbone/side-chain. Although the peptide chain becomes more compact at short N-C termini distances, this structural transformation is manifested much more in the solvent exposure of the side-chain than in the backbone.

**Figure 6:**
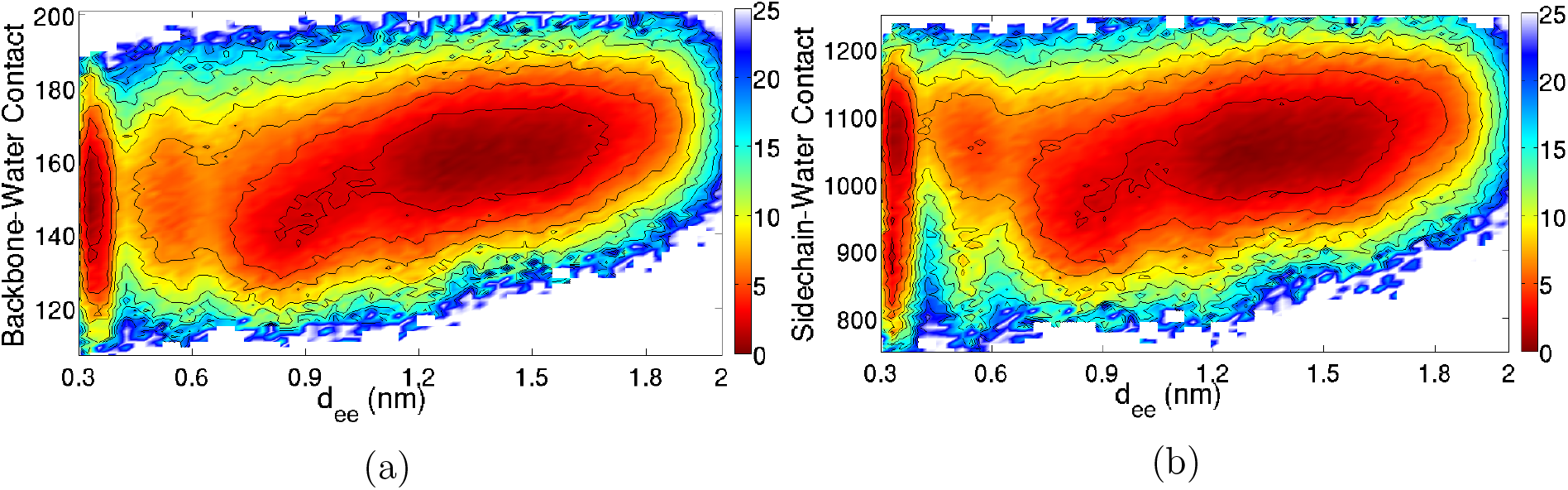
Reweighted 2D FES along *d*_*ee*_ and backbone-water contact(a) and along *d*_*ee*_ and sidechain-water contact(b)

In the following section, we will tackle the exposure of the peptide to the solvent through an examination of some topological properties involving the hydrogen bond network. The snapshots shown earlier in Figure 5 depicts the physical origins of Figure 6. In one case (panel b) the side chains are mostly separated away from each other leaving a lot of room for water molecules to be interspersed between the side chains. On the other hand, for state A which is the most compact state, involves the collective collapse of several of these side chains leading to the expulsion of solvent. The backbone on the other hand, as we will see in more detail in the next section, still remains solvent exposed - in fact, in both state A and B we see that the amide dipoles orient in such a fashion so that several N-H groups of the backbone essentially form strong hydrogen bonds with a single C=O, while most of the other carbonyl oxygens remain solvent exposed. The fact that the anionic carbonyl groups are exposed to water is consistent with an idea proposed by Collins^73^ who suggested that the structuring of water induced by ions, is rooted in specific hydrogen bonding interactions. He proposed that anions are better at ordering water molecules than cations due to the asymmetry of charge in a water molecule. In the next section we will see how the ordering of water is reflected in the formation of water wires around the peptide chain.

### 3.2 Water Wires Around Amyloid *β*_30–35_ Chain

Earlier we showed that the backbone-water contact map does not feature any significant changes during the structural conformational changes. On the other hand, there are much larger changes in the solvent density around the hydrophobic side-chains. The backbone-water contact map quantifies the polar interactions formed between the amide groups and the surrounding water but is blind to the orientational correlations originating directly from the hydrogen bond network.

Understanding the role of hydrogen bond networks in shaping both the structural and dynamical properties of biological systems continues to be a topic of active investigation.^74–77^ Thirumalai and co-workers have also shown that long-lived water wires between two beta sheets from the polar fragment of the yeast prion protein, result in long-lived metastable structures.^47,48^ The analysis of the water wires in these studies is done in a qualitative manner without quantitatively probing the hydrogen bond network. There have been several theoretical studies examining the importance of water networks in maintaining the structural integrity of proteins as well as in the interaction of proteins. In particular, Mazen A. and co-workers^77^ showed that during the encounter of two hydrophilic proteins, adhesive water networks form between them stabilizing intermediate states before native contacts form. In all these studies, the connections between the nodes in the network neglect the directionality of the interactions. It is clear however, that if one is interested in networks associated with water hydrogen bonds as we are here, then one must consider the evolution of directed networks.

In the following section we examine the cooperative and collective behavior associated with the reorganization of the directed adhesive networks enveloping the hexapeptide chain in water. As described in the Methods section, we examined the statistics associated with the shortest directed path connecting candidate donor and acceptor groups. Before we showcase individual water wires forming an important and integral part of the protein structure and how they change during the conformational fluctuations, we begin by quantifying how the collective directed network from all possible donor to acceptor groups evolves during conformation fluctuations of protein. In order to quantify this, we examined the global efficiency of the protein-water network which is defined as^78–80^

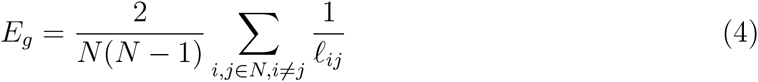

where N is the number of vertices (N-H and C=O groups) and ℓ_*ij*_ is the shortest path length between i and j vertices. If the two vertices i an j are not connected, 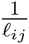 is set to zero so that these pairs don't contribute to *E*_*g*_.

Figure 7 shows the FES obtained along *d*_*ee*_ and *E*_*g*_. Since the global efficiency averages over many directed paths from different parts of the peptide, it is useful to calibrate ourselves first on the typical shortest path lengths (*l*_*ij*_) that the *E*_*g*_ maps onto. An average directed path of length 1 would correspond to an *E*_*g*_ of 1. In the limit that there are no directed paths between the nodes, *E*_*g*_ would tend to 0. When the N and C termini are separated far away from each other, Figure 7 shows that *E*_*g*_ is peaked at 0.065 corresponding to an average path length of about 16. In this extended state, there are large fluctuations in the directed network connecting the donor N-H and C=O groups in the peptide as seen in the broad distribution of *E*_*g*_ extending from 0.04 to 0.125 implying the formation of very long wires of length 25 to rather short wires of length 8. Moving from states C and D into the regime where the N and C termini are in close contact, leads to an overall increase in *E*_*g*_. At short *d*_*ee*_ distances of 0.35 nm, there is a minimum corresponding to a typical path length of 11 which result from the formation of shorter and directed paths between the donor and acceptor groups compared to the extended state although there are still rather large fluctuations in the network.

**Figure 7:**
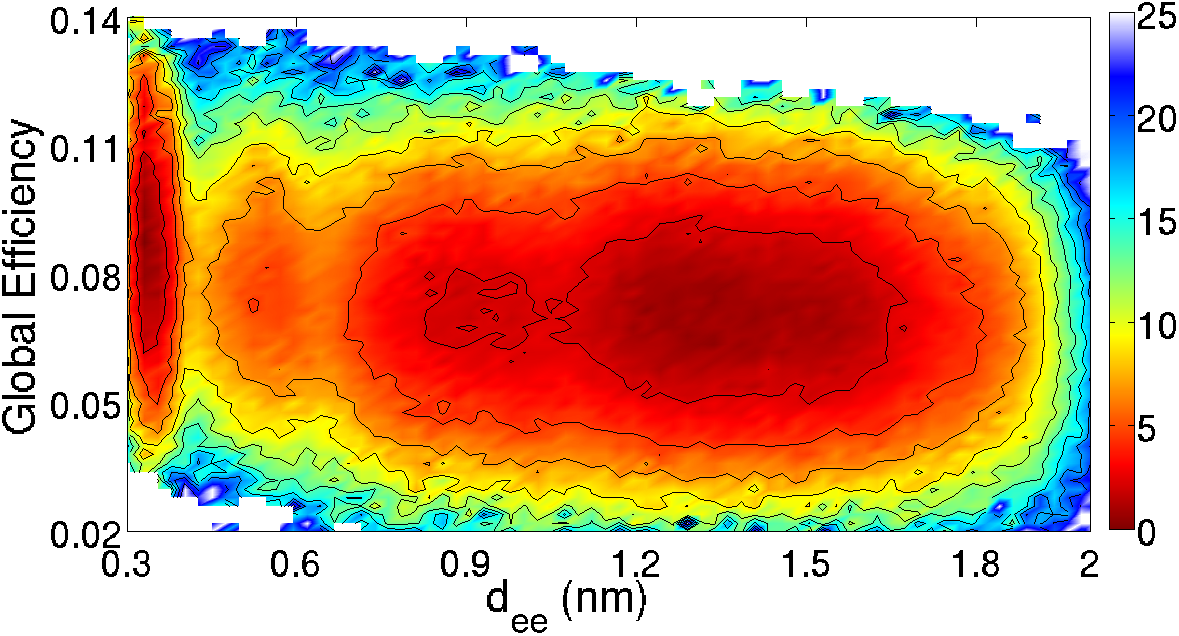
Reweighted 2D FES along *d*_*ee*_ and global efficiency

The global efficiency of the network averages over connections involving all terminus-terminus, termini-backbone and backbone-backbone and hence we get a global measure of the changes in the connectivity. Furthermore, the contributions to *E*_*g*_ between candidate donor and acceptor groups involve the possibility of *many* long wires or *a few* short wires. To understand better the changes of the water networks involving the termini and backbone separately, we show in Figure 8 the FES obtained by examining the components of local network. The top left panel elucidates the changes involving the specific connections between the N and C terminus - the transition from the extended to contact pair involving the formation of a strong salt bridge. This salt bridge consists of both direct contact between the N and C termini as well as by water mediated interactions due to the formation of short water wires ranging from length 2 to 5. Examples of the type of water wires are shown in the bottom left panel of Figure 8.

**Figure 8:**
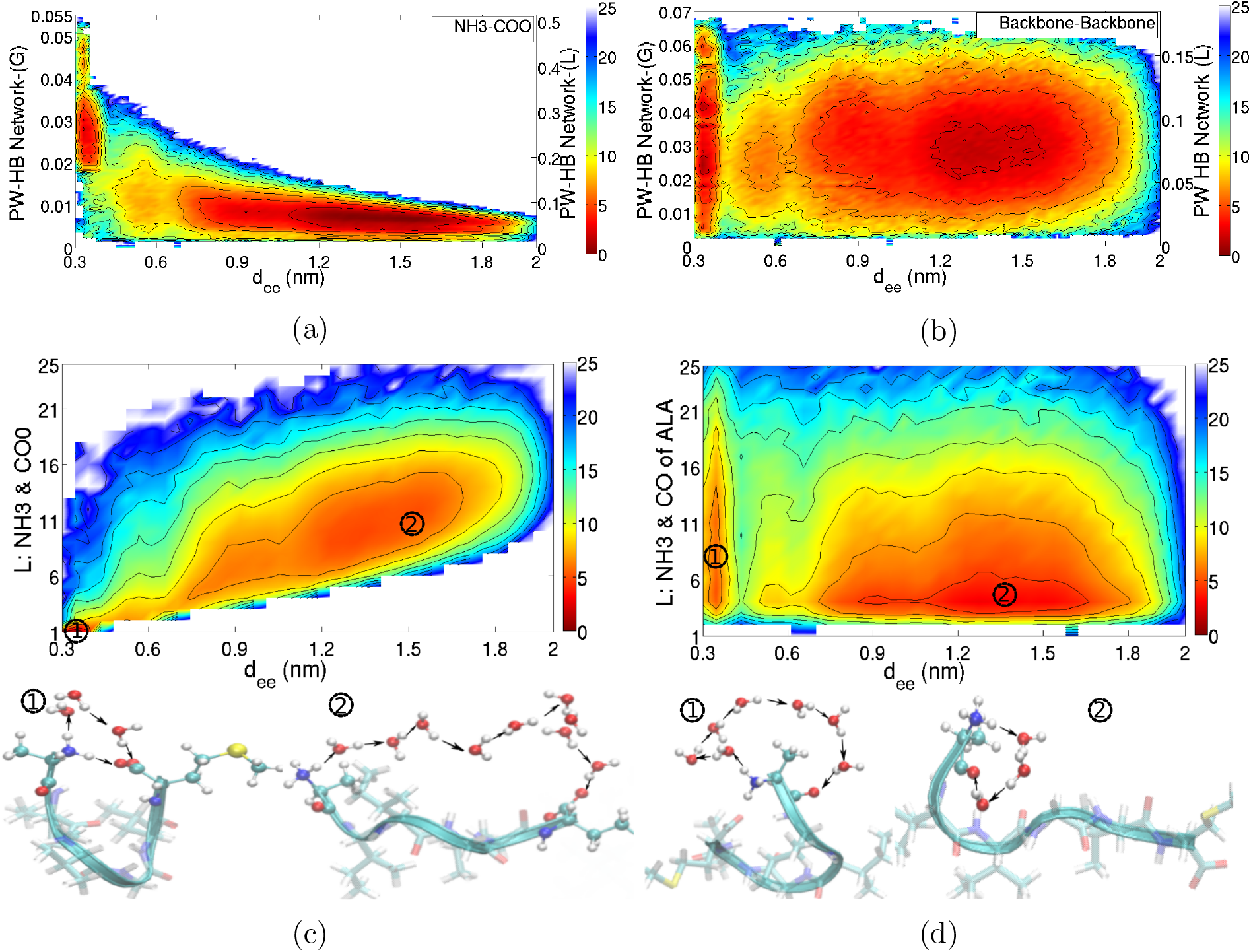
Top: reweighted 2D FES along *d*_*ee*_ and protein-water H-bond (PW-HB) network connectivity between termini NH3 and COO(a), PW-HB along *d*_*ee*_ and protein-water network connectivity between NH and CO of backbone (b). Bottom: reweighted 2D FES along *d*_*ee*_ and L of NH3-COO (c) and along *d*_*ee*_ and L of NH3-CO(ALA)(d). L indicated here is an equivalent short notation to ℓ_*ij*_ in Eq. 4

The top right panel of Figure 8 elucidates the water networks formed between all the backbone amide groups and CO. In the extended state, the backbone is stabilized by water wires of varying lengths ranging from 5 to 30 with a dominant peak at 15. Transitioning into the regime where the N and C termini are in close contact results in a more structured landscape implying that in both states A and B, there are different types of sub-states stabilized by unique water-networks. Some examples of these water networks are illustrated in the bottom right panel of Figure 8. A similar type of analysis focusing on the networks connecting the N-terminus to the backbone and the backbone to the C-terminus shows distinct patterns in the water wires between the two respective situations. To illustrate these features we show in the left and right panels of Figure 9 we compare the distribution of water wires from the N-terminus to the C=O group of Glycine and from the N-H group of the Glycine to C-terminus respectively. The free energy landscapes associated with these coordinates are rather distinct reinforcing the notion that the interactions of the N and C termini with the backbone is mediated by specific water-wire motifs. For more details on the global and local-efficiency network measures associated with the NH_3_(terminus)-backbone and backbone-COO^−^(terminus), the reader is referred to the SI (Sect. SI-5).

**Figure 9:**
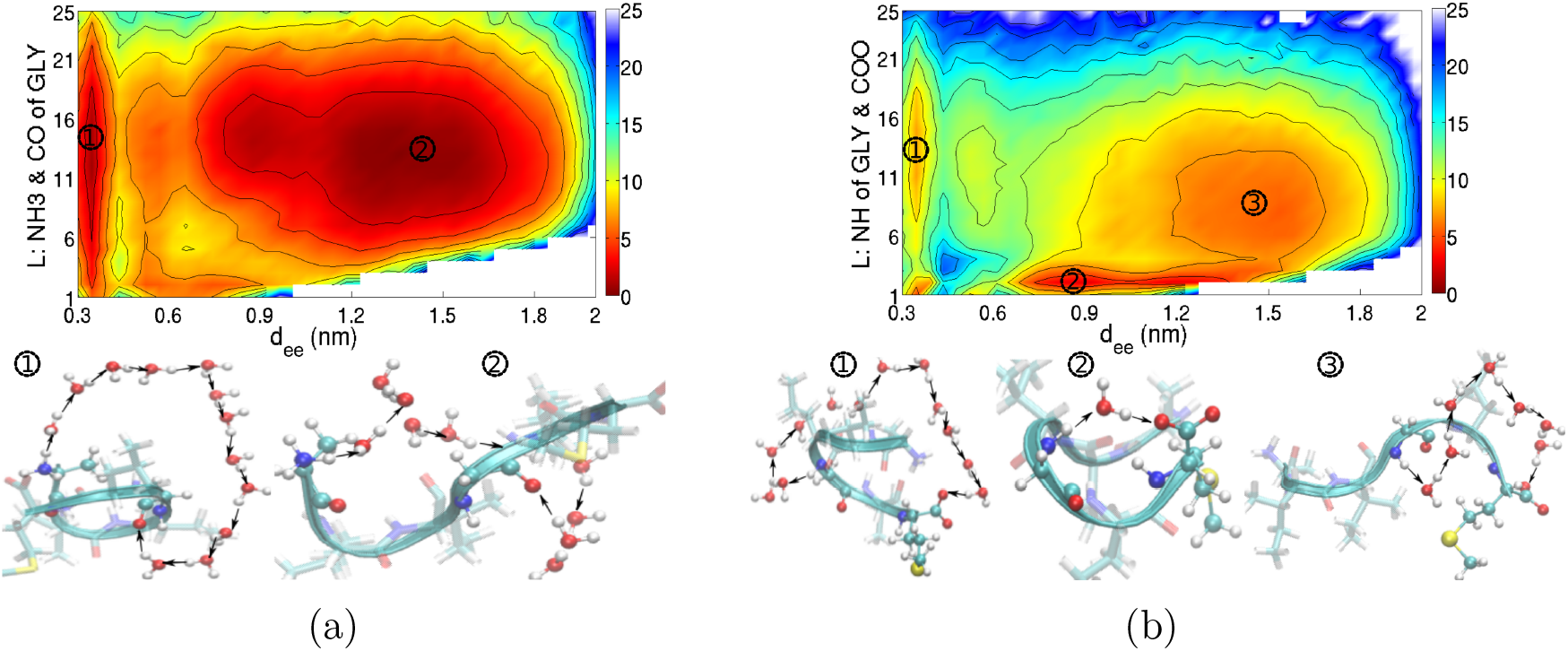
Reweighted 2D FES along *d*_*ee*_ and L of NH3-CO(GLY) (a) and along *d*_*ee*_ and L of NH(GLY)-COO (b)

## 4 Discussion and Conclusion

In this work, we have explored the conformational landscape of a hexapeptide of the C-terminal fragment of Amyloid *β*_30–35_ in liquid water using well-tempered metadynamics simulations. The free energy landscape of this system is very rich and characterized by an underlying structural disorder where the N and C termini plays a critical role in stabilizing different conformations. The conformational fluctuations on the free energy landscape is driven by an intimate coupling of pure electrostatic, hydrogen bond networks and hydrophobic forces. Although the importance of these interactions are known to be important in the aggregation of biological systems, a molecular characterization and understanding of the underlying processes still remains an active area of research.

Here we placed particular emphasis on the coupled reorganization of the hydrogen bond networks involving intraprotein contacts as well as that of the protein and surrounding water. The electrostatic interaction between the N and C termini results in the formation of strong salt-bridges that can be both in direct contact or mediated through the solvent by water wires. Besides the salt bridges, there are a myriad of network connections that can form between different donor and acceptor groups of the peptide through the solvent. Thus the different conformations that can be stabilized are best seen as a collective reorganization of both protein and water networks. At the same time, the side chains of the amino acids are hydrophobic in nature and the extent to which they pack closely with each other, also plays an important role in stabilizing different types of conformations. The collapse of the sidechains and explusion of water results in the formation of a protein conformation resembling a Janus-like particle.

The coupling of the reorganization of the water networks as well as the collapse of the hydrophobic side-chains, as pointed out earlier, is shown more clearly in the left and right panels of Figure 10. The left panel shows the coupling between the *squeeze out* of water molecules between the side-chains and the increase in their van-der-Waals packing. The transition from states B to A involves a coupling of these two events occurring simultaneously similar to what was observed in earlier simulations of a hydrophobic sub-sequence of the amyloid fibril.^48^ Besides the water explusion of water, there is also a collective reorganization of the hydrogen bond network. This feature is seen in the right panel of Figure 10 where we see that during the collapse of the hydrophobic side-chains there is an overall shortening of water wires enveloping the hexapeptide. It would be interesting in future studies to understand how these features change as one moves to larger polypeptides as well as how the mechanisms are altered when the chemistry of the side-chain groups are altered.

**Figure 10:**
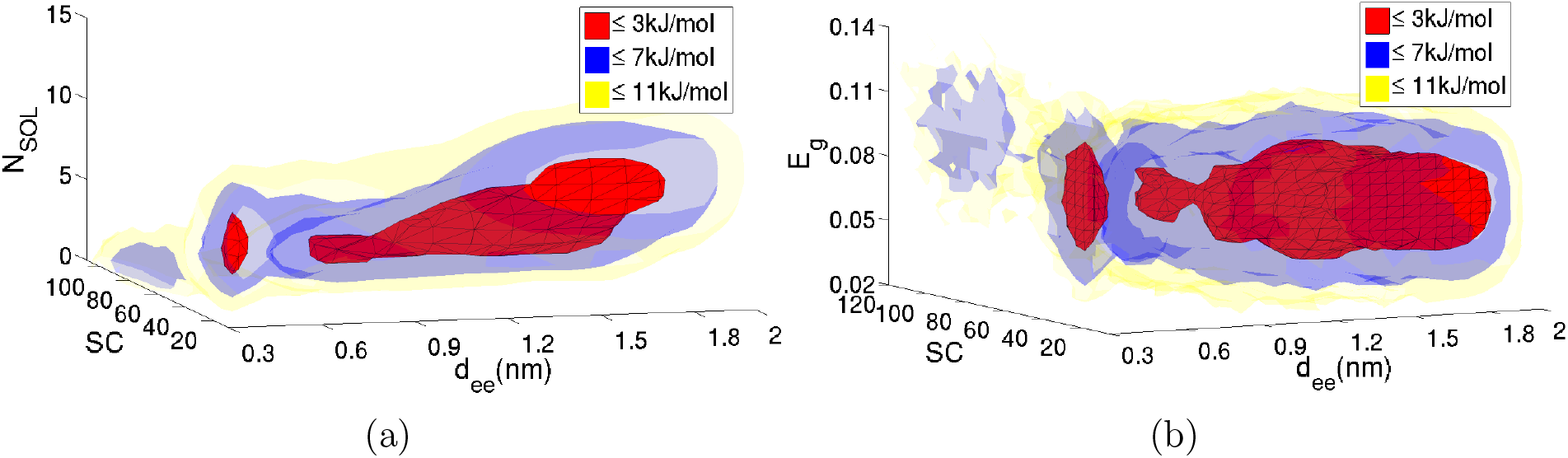
Reweighted 3D FES along *d*_*ee*_, *R*_*g*_ and number of water molecules inside sidechain shell of radius 0.5nm(a) and along *d*_*ee*_, sidechain contact and global efficiency(b)

It is clear that the the N and C termini play an important role in stabilizing different conformations of the hexapeptide in water. These conformations involve explicit salt-bridges between the termini and strong hydrogen bonds between the termini and the backbone amide groups. As indicated earlier, in order to assess the sensitivity and to understand how the free energy landscape changes in the absence of standard N and C terminal groups, we also performed simulations of a hexapeptide that is capped with methyl molecules. The absence of the standard termini groups has a rather drastic effect on the conformational landscape. In Figure 11, the FES along both *d*_*ee*_ and *R*_*g*_ confirms the presence of essentially only one minima. Interestingly, if we compare this to the FES of the NC system, they share similar features of having analagous states A through D. However, the absence of the termini interactions de-stabilizes many of the conformations and hence is likely to affect the mechanisms associated with amyloid aggregation inferred in previous simulations where the termini are capped with methyl groups^20,81–84^

**Figure 11:**
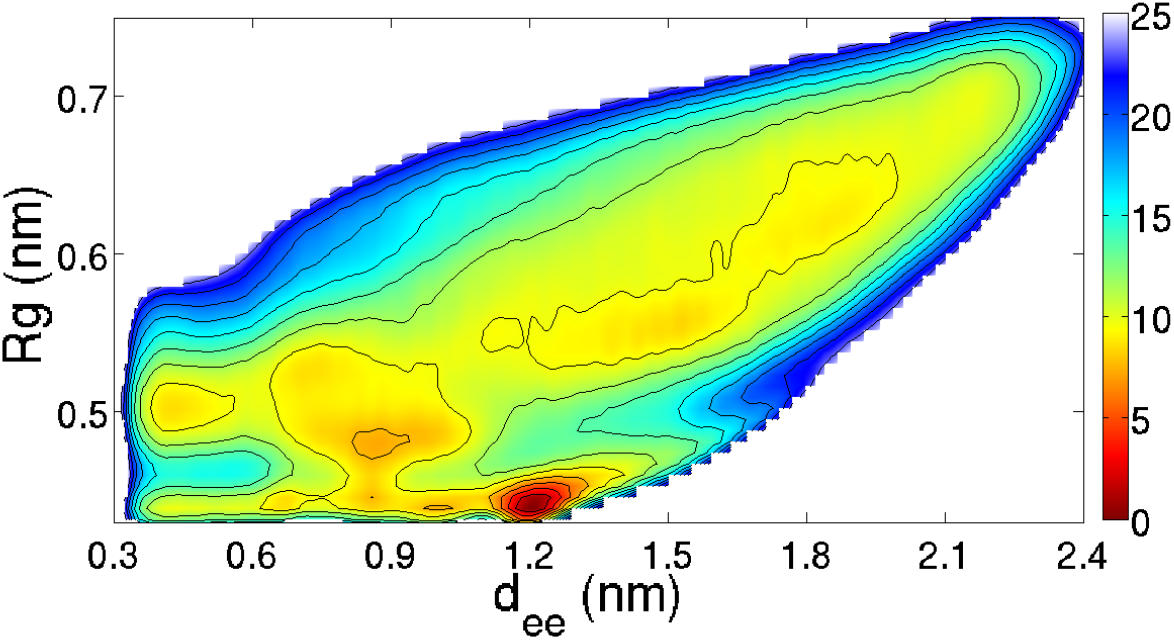
2D FES obtained from 1.1-*μ*s-long WT-MetaD for MET chain. The FES is contoured by 2.5kJ/mol up to 25kJ/mol.

With the computational methodology applied in this work, based on WT-MetaD, recovering time-correlation information is not possible. This will allow to directly explore interesting time-dependent quantities such as frailty of H-bond networks, transition times between different basins and causality in conformational changes. In spite of being straight-forward, in the future we will investigate the possibility to extract this information from current dynamics, possibly using some already available theoretical tools.

Although our simulations only examine the importance of the evolution and fluctuation of directed water wires for the monomer chain, it would be interesting to explore in the future, how these properties change during the aggregation process and how this is altered by the number of monomers in the nucleation center.^47^ Our studies should also motivate new experiments examining the effect of mutating the termini on spectroscopic observables probed through methods such as circular dichroism^85^ and nuclear magnetic resonance^86^ and also on optical properties such as fluorescence.^34^

## 5 Extra information when writing JACS Communications

### Acknowledgement

#### Supporting Information Available

This material is available free of charge via the Internet at http://pubs.acs.org/.

